# Chronic estrus disrupts uterine gland development and homeostasis

**DOI:** 10.1101/347757

**Authors:** C. Allison Stewart, M. David Stewart, Ying Wang, Rui Liang, Yu Liu, Richard R. Behringer

**Affiliations:** department of Genetics, University of Texas M.D. Anderson Cancer Center, Houston, TX, USA; Department of Biology and Biochemistry, University of Houston, Houston, TX, USA

**Keywords:** GnRHR, progesterone, endometrium, adenogenesis, neoplasia

## Abstract

Female mice homozygous for an engineered *Gnrhr* E90K mutation have reduced gonadotropin-releasing hormone signaling, leading to infertility. Their ovaries have numerous antral follicles but no corpora lutea, indicating a block to ovulation. These mutants have high levels of circulating estradiol and low progesterone, indicating a state of persistent estrus. This mouse model provided a unique opportunity to examine the lack of cyclic levels of ovarian hormones on uterine gland biology. Although uterine gland development appeared similar to controls during prepubertal development, it was compromised during adolescence in the mutants. By 20 weeks of age, uterine gland development was comparable to controls, but pathologies, including squamous neoplasia, tubal neoplasia, and cribriform glandular structures, were observed. Induction of ovulations by periodic human chorionic gonadotropin treatment did not rescue post-pubertal uterine gland development. Interestingly, progesterone receptor knockout mice, which lack progesterone signaling, also have defects in post-pubertal uterine gland development. However, progesterone treatment did not rescue post-pubertal uterine gland development. These studies indicate that chronically elevated levels of estradiol with low progesterone and therefore an absence of cyclic ovarian hormone secretion disrupts post-pubertal uterine gland development and homeostasis.

## Introduction

Endometrial or uterine glands produce and transport secretions during gestation that are essential for embryo survival and development (Gray et al., 2001). This is especially true during pre-implantation development, when uterine glands are the primary source of nutrients for the embryo. The uterine secretome has been characterized and includes nutrients, cytokines, hormones, and a variety of other factors (Bhutada et al., 2013; Bhusane et al., 2016; Salamonsen et al., 2016). Blocking uterine gland formation causes infertility due to failed implantation, highlighting the essential role for uterine glands in early conceptus development (Gray et al., 2001; Gray et al., 2002; Dunlap et al., 2011).

Uterine gland development proceeds in two waves. In most species studied, uterine glands are not present at birth when the uterus consists of a simple epithelium surrounded by undifferentiated mesenchyme (Hu et al., 2004; Cooke et al., 2013). Shortly after birth, the uterus differentiates into separate layers, an endometrial stromal layer surrounded by two myometrial smooth muscle layers (Hu et al., 2004). At birth, the endometrium consists of a simple, columnar epithelium, lining the lumen of the uterus (luminal epithelium) and the underlying stroma. In the mouse, the first wave of uterine gland development occurs within a week after birth, when numerous foci within the luminal epithelium begin to invaginate and elongate through the stroma toward the myometrium (Cooke et al., 2012; Vue et al., 2018). During this prepubertal period, uterine gland development occurs in an ovary- and hormone-independent manner (Branham and Sheehan, 1995; Ogasawara et al., 1983). The second wave of uterine gland development occurs after puberty (adolescence) and is dependent upon ovarian hormones (Stewart et al., 2011). At this stage, uterine gland growth elaborates branched structures within the stroma. In primates and a subset other mammalian species, a third period of growth occurs with every menstrual cycle as the endometrium is regenerated (Maybin and Chritchley, 2011; Rasweiler, 1991; van der Horst and Gillman, 1941).

The gonadotropin-releasing hormone receptor (GnRHR) is a seven-transmembrane G-protein coupled receptor primarily expressed by gonadotroph cells in the anterior pituitary (Stamatiades and Kaiser, 2017). GnRHR signaling stimulates the release of the gonadotropins, luteinizing hormone (LH) and follicle-stimulating hormone (FSH) in response to its ligand, GnRH produced in the hypothalamus. The pulsatile release of LH and FSH is essential for ovarian follicle selection, maturation, ovulation and luteinization, and production of the ovarian steroid hormones, estradiol and progesterone. Steroid hormones form a negative feedback loop for GnRH release. This system is referred to as the hypothalamic-pituitary-gonadal axis, which is essential for cyclicity, fertility, and pregnancy. GnRHR signaling stimulation by GnRH agonists has been used to treat disorders in women, including endometriosis, uterine fibroids, and polycystic ovarian syndrome (PCOS) (Hackenburg et al., 1992; Marschalek et al., 2015; Tsai et al., 2017). However, GnRH agonist treatment can increase the risk of pathologies, including diabetes mellitis and cardiovascular disease (Palomba et al., 2004). Inhibition of GnRH signaling by GnRH antagonists has been used during ovarian hyperstimulation for fertility treatments (Lee et al., 2008). GnRH antagonist treatment can prevent the development of diabetes in susceptible rodents (Ansari et al., 2004). It has also been used to reduce testosterone levels in prostate cancer patients (Steinberg, 2009). Thus, modulating GnRH signaling regulates a variety of physiological processes and can lead to pathologies.

Previously, we engineered mice with a single amino acid substitution in the GnRHR gene (*Gnrhr* E90K) to mimic a human homozygous mutation that causes hypogonadotropic hypogonadism in males (Söderlund et al., 2004; Stewart et al., 2012). The E90K substitution prevents formation of a salt bridge between E90 and K121 in the first and second transmembrane domains (Janovick et al., 2007). This folding defect is recognized and retained in the endoplasmic reticulum by the quality control system. Interestingly, if correctly routed to the plasma membrane, the E90K mutant responds to GnRH and functions appropriately (Janovick et al., 2013). The net result of the E90K mutant is reduced plasma membrane expression of GnRHR and therefore reduced signaling. *Gnrhr*^*E90K*^ homozygous female mice are infertile due to a block in ovulation. Their ovaries have multiple, antral follicles but no corpora lutea (CL) (Stewart et al., 2012). Thus, *Gnrhr*^*E90K*^ homozygous females do not cycle, suggesting that they are in a persistent state of estrus. These mice provided a unique model to determine the consequence of preventing cyclic ovarian hormone production on uterine gland development. In these mutant mice, post-pubertal uterine gland development was impaired and disease-related phenotypes occurred, including uterine edema and endometrial neoplasia. These results suggest that cyclic steroid hormone production is essential for post-pubertal uterine gland development and endometrial homeostasis.

## Materials and Methods

### Mice

The *Gnrhr*^*E90K*^ and *pgr*^*tm2(cre)Lyd*^ mice were maintained on a C57BL/6J × 129/SvEv mixed background. *Gnrhr*^*E90K*^ and *pgr*^*tm2(cre)Lyd*^ mice were genotyped as previously described (Soyal et al., 2005; Stewart et al., 2012). Mice for experimental analysis were obtained by crossing *Gnrhr*^*E90K*/+^ females with *Gnrhr*^*E90K/E90K*^ males. The mutants were *Gnrhr*^*E90K/E90K*^ and the controls were *Gnrhr*^*E90K*/+^. For *pgr*^*tm2(cre)Lyd*^ (*Pgr*^*Cre*^) mice, *Pgr*^*Cre*/+^ females were bred with *pgr*^*Cre/Cre*^ males. The mutants were *pgr*^*Cre/Cre*^ (referred to as PRKO) and the controls were *Pgr*^*Cre*/+^ (referred to as controls) All experimental procedures involving mice were approved by the Institutional Animal Care and Use Committee of the M.D. Anderson Cancer Center.

### Hormone treatments

Human chorionic gonadotropin (Sigma) or PBS was injected intraperitoneally every 4 days from 5 to 8 weeks of age. Slow-release implants were prepared with or without 0.5 g/ml progesterone in peanut oil in silastic tubing (Fisher Scientific 11-189-15G) with a 1 cm release space. The ends of the implants were heat sealed with polyethylene tubing inserted into the silastic tube (Cohen and Milligan 1993; Milligan and Cohen 1994; Yellon et al., 2009).

### Tissue collection

Tissues were dissected in phosphate buffered saline (PBS) immediately after euthanasia. Uterine horn length was measured from the uterotubal junction to the uterine bifurcation on freshly dissected tissues. The number of CL were counted in freshly dissected ovaries. Uteri and ovaries were fixed in 4% paraformaldehyde in PBS for 24 h at 4°C and rinsed in 70% ethanol for ≥ 24 h prior to tissue processing. All tissues were dehydrated through a standard ethanol gradient, cleared in Histoclear (National Diagnostics) and embedded in Paraplast Plus^®^ Tissue Embedding Medium (Statlab).

### Histological analyses and glandular tissue quantification

Five μm sections were cut from paraffin embedded tissues. H&E staining was performed using a standard protocol. Uterine glandular tissue abundance was performed by counting the number of endometrial gland cross-sections (5 slides, 6 sections per slide for each animal) of the uterus from mice of each genotype at various time points or following hormone treatment (n = 4-9) in a blind experiment.

### Microscopy

Gross images were acquired using a Leica MZ10F stereomicroscope and the extended depth of focus feature of the LAS v3.7 software (Leica Microsystems, Wetzlar, Germany). Brightfield images of tissues sections were obtained using a Nikon 80i microscope equipped with a DS-Fi1 5-megapixel color camera (Nikon Instruments) and NIS Elements AR 3.1 software v4.13 (Nikon Instruments).

### Real-time PCR

Total RNA for quantitative (Q)PCR was extracted using TRIzol reagent according to the manufacturer’s protocol (Life Technologies, Carlsbad, CA). Relative mRNA levels were measured using TaqMan gene expression assays with 6-carboxyfluorescein (FAM)-labelled probes. Primer probe sets were: *Cyp19a1* (Mm00484049(m1), *Cyp11a1* (Mm00490735_m1), *Esr1* (Mm00433149_m1), *Pgr* (Mm00435625_m1), *Foxa2* (Mm00839704_mH), *Ihh* (Mm00439613_m1). *Gapdh* mRNA expression levels were used for normalization. The PCR was run using an ABI Prism 7900HT thermocycler and SDS2.1 software (Applied Biosystems). Data were analyzed by the comparative ΔΔCT method.

### Serum hormone levels

Serum levels of progesterone, estrogen, and testosterone were measured by ELISA by the Ligand Assay and Analysis Core at the University of Virginia (Charlottesville).

### Statistical analyses

For experiments with more than two experimental groups, quantitative data were subjected to one-way analysis of variance (ANOVA) using MedCalc v14.12.0 software (MedCalc Software bvba, Belgium). If the ANOVA was positive (*P* < 0.05), a post-hoc Student-Newman-Keuls test was performed for pairwise comparison of subgroups. For studies with two experimental groups, an independent samples Student’s *t*-test was performed. A *P*-value of < 0.05 was considered significant for all tests.

## Results

### Ovulation failure in *Gnrhr^E90K^* homozygotes results in persistent estrus and low levels of progesterone

Previously, we reported that female mice homozygous for *Gnrhr*^*E90K*^ are infertile due to ovulation failure (Stewart et al., 2012). In this study, our objective was to determine how ovulation failure affected cyclic hormone levels and uterine gland development. At 8 weeks of age, female mice heterozygous for *Gnrhr*^*E90K*^ (controls) were fertile and cycled. Ovarian histology showed various stages of folliculogenesis with multiple CL (**Fig. 1A**). At estrus, *Gnrhr*^*E90K*^ heterozygotes exhibited multiple antral follicles and CL (**Fig. 1B**). In contrast, *Gnrhr*^E90K^ homozygous females had multiple antral follicles with no CL (**Fig. 1C**). Similar results were found at 20 weeks of age. *Gnrhr*^*E90K*^ heterozygotes exhibited a mixture of follicles at various stages and CL (**Fig. 1D**). However, no CL were observed on the ovaries of *Gnrhr*^*E90K*^ homozygotes (**Fig. 1E**). The average number of CL per mouse (pair of ovaries) is summarized in **Fig. 1F**.

We next determined the effect of the GnRHR E90K-induced ovulation block on blood levels of ovarian steroid hormones. Serum was collected at estrus for all groups to normalize for the persistent estrus observed in *Gnrhr*^*E90K*^ homozygotes. At 3 weeks of age, progesterone levels were comparable between mutants and controls (**Fig. 1G**, P=0.29). As expected due to the lack of CL, progesterone levels were reduced at 8 weeks of age (**Fig. 1G**, P<0.05). Estradiol levels were similar between mutants and controls at 3, 8, and 20 weeks of age (**Fig. 1H**, P=0.21, P=0.32, P=0.1, respectively) Testosterone levels were similar between mutants and controls at 8 and 20 weeks of age (**Fig. 1I**, P=0.41, P=0.24, respectively). These data support the idea that *Gnrhr*^*E90K*^ homozygotes exhibit persistent estrus with high estradiol and low progesterone levels.

**Fig. 1.**
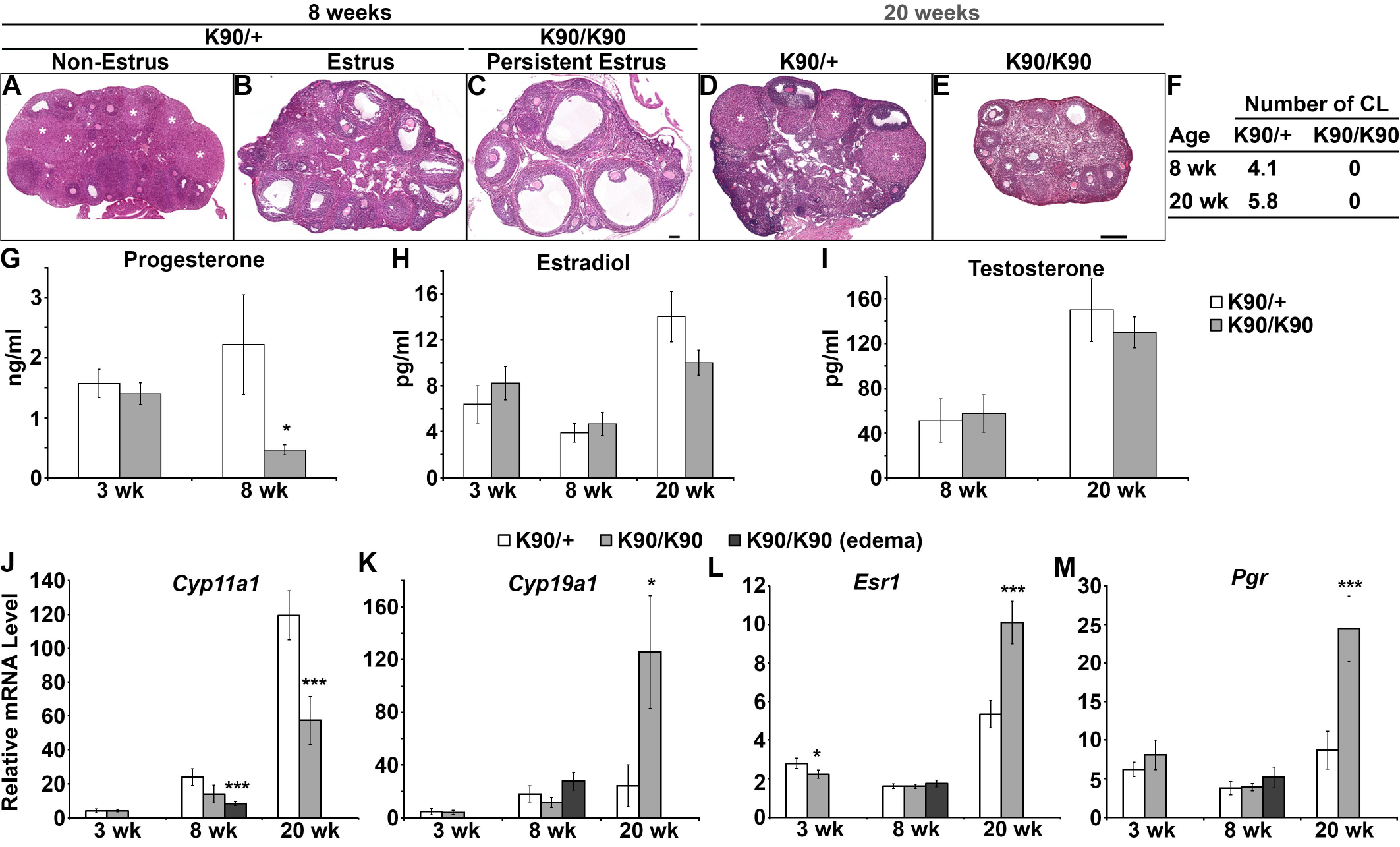
Ovulation failure in *Gnrhr*^*E90K*^ homozygotes results in persistent estrus and low levels of progesterone. (A-E) H&E-stained ovarian sections. Corpora lutea (CL) (white asterisks) were abundant in heterozygous controls (A, B, D), but absent in *Gnrhr*^*E90K*^ homozygotes (C, E). A-C, scale bar = 100μm; D, E, scale bar = 250 μm. (F) Average number of CL per ovary (± SEM). (G-I) Steroid hormone levels. (J-M) Realtime PCR gene expression levels (± SEM). * P<0.05, *** P < 0.005, Student’s t-test. K90/+, *Gnrhr*^E90K^ heterozygotes; K90/K90, *Gnrhr*^*E90K*^ homozygotes.

Next, we tested the effect of GnRHR E90K on the expression of genes encoding steroidogenic enzymes (*Cyp11a1* and *Cyp19a1*, encoding P450scc and aromatase, respectively) and steroid hormone receptors (*Esr1* and *Pgr*, encoding estrogen receptor alpha and progesterone receptor, respectively) in the ovary. P450scc catalyzes the first biosynthetic step in steroidogenesis, converting cholesterol to pregnenolone. Aromatase converts testosterone to estradiol.

At 3 weeks of age, no differences in the levels of *Cyp11a1* (P=0.47), *Cyp19a1* (P=0.42), or *Pgr* (P=0.18) were observed between mutants and controls; however, *Esr1* was reduced in the mutants (P<0.001). At 8 and 20 weeks of age, *Cyp11a1* was down-regulated in mutants compared to controls (**Fig. 1J, K**, P<0.005 for both time points). *Cyp19a1* was up-regulated at 20 weeks of age (**Fig. 1J, K**, P=0.01). At 8 weeks of age, *Esr1* (P=0.48) and *Pgr* (P=0.44) expression levels were comparable between the mutants and controls but were elevated at 20 weeks of age (**Fig. 1L, M**, P<0.001, P<0.002, respectively).

### Chronic estrus results in compromised uterine gland development during adolescence

Uterine gland development proceeds in two waves. The first glands form as buds from the luminal epithelium shortly after birth (Hu et al., 2004; Vue et al., 2018). This initial gland structure is elaborated during adolescence. Postnatal uterine gland development in *Gnrhr*^E90K^ homozygous females was similar to controls as determined by histology (**Fig. 2A-D**), measurements of uterine horn length (**Fig. 2M**, P=0.41) and glandular tissue abundance (**Fig. 2N**, P=0.23). By 8 weeks of age (post-puberty), uterine histoarchitecture was affected in *Gnrhr*^*E90K*^ homozygotes. About half of the *Gnrhr*^*E90K*^ homozygotes exhibited uteri that appeared similar to controls except with less uterine glandular tissue abundance (**Fig. 2F, I**). The other half the mutants had severe uterine edema accompanied by a reduction in myometrial and endometrial layer width and very few uterine glands (**Fig. 2G, J**). At 8 weeks of age, uterine horn length was increased for *Gnrhr*^*E90K*^ homozygotes compared to controls (**Fig. 2M**, P<0.001). In addition, uterine glandular tissue abundance was significantly decreased (**Fig. 2N**, P<0.001). These data demonstrate impaired uterine gland development during adolescence in *Gnrhr*^*E90K*^ homozygous females that coincides with loss of cyclic steroid hormone production.

**Fig. 2.**
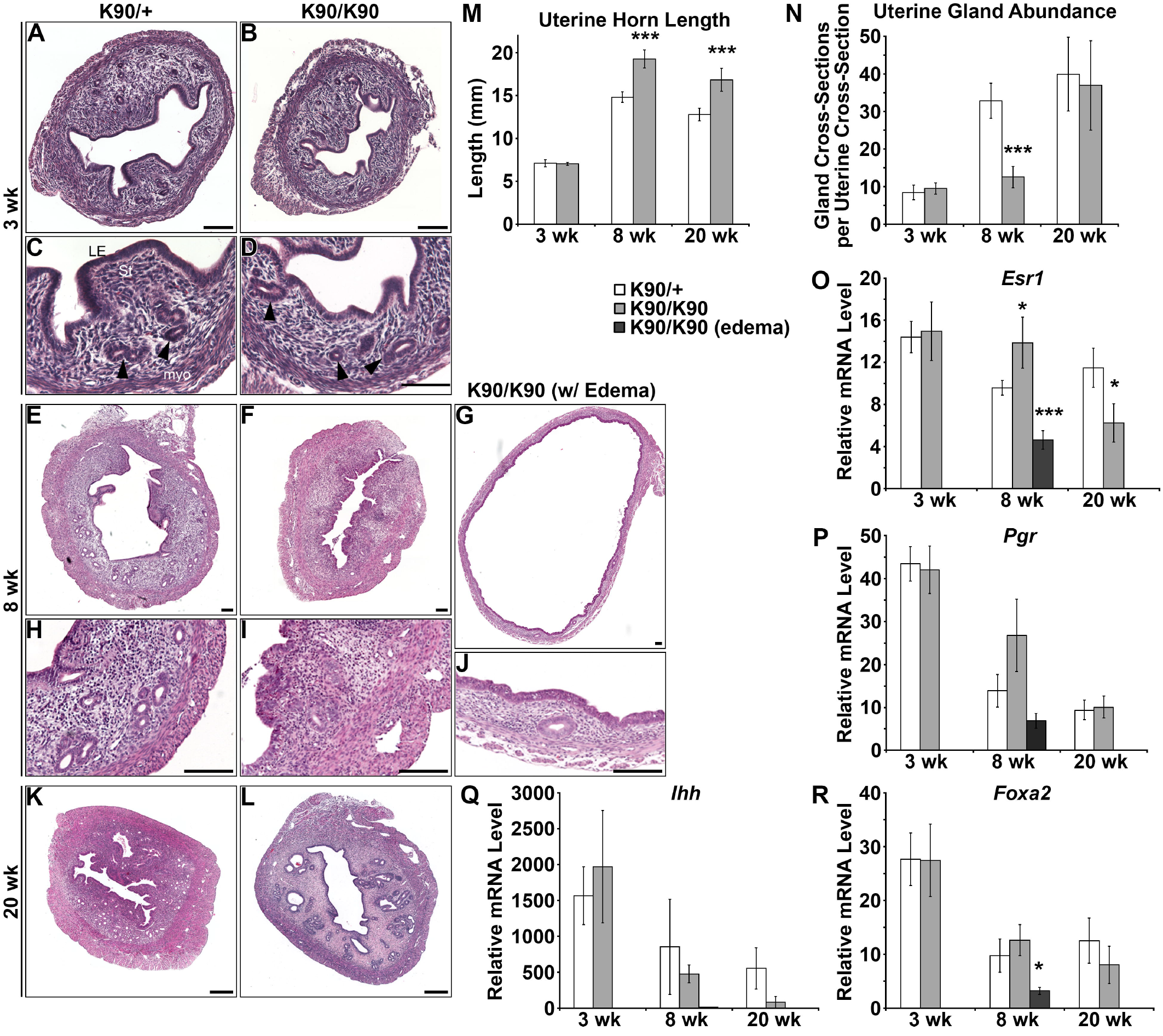
Compromised uterine gland development during adolescence. (A-L) H&E-stained uterine sections. (A, C, E, H, K) *Gnrhr*^*E90K*^ heterozygous controls. (B, D, F, G, I, J, L) *Gnrhr*^*E90K*^ homozygous mutants. Uterine histoarchitecture and gland development was comparable between mutants and controls at 3 weeks of age (A-D). Uterine gland development was perturbed in the mutants by 8 weeks of age (E-J). Some *Gnrhr*^*E90K*^ homozygotes exhibited uterine edema (G, J). By 20 weeks of age, uterine gland development was similar between mutants and controls (K, L). Scale bars = 100 [jm. Black arrowheads, uterine gland tissue. (M) *Gnrhr*^*E90K*^ homozygotes displayed increased uterine horn length after 3 weeks of age. (N) The mutants had decreased numbers of uterine glands at 8 weeks of age. (O-R) Expression levels of *Esr1*, *Pgr*, *Ihh*, and *Foxa2*. * P<0.05, *** P < 0.005, Student’s *t*-test.

Interestingly, the uteri of *Gnrhr*^*E90K*^ homozygous females at 20 weeks of age were similar to controls. No uterine edema was detected and uterine histoarchitecture was comparable to controls (**Fig. 2K, L**). Uterine horn length was still increased in the mutants compared to controls (**Fig. 2M**, P<0.001), but uterine glandular tissue abundance was similar to controls (**Fig. 2N**, P=0.27). However, pathologies were observed (see below). These data suggest the post-pubertal wave of uterine gland development is delayed in the absence of cyclic steroid hormone secretion.

At 3 weeks of age, uterine expression of *Esr1* (P=0.42) and *Pgr* (P=0.41) was similar, consistent with histological assays. At 8 weeks of age, *Esr1* mRNA levels were elevated in non-edematous uteri of mutants (P=0.04), but decreased in edematous mutant uteri relative to controls (P<0.001). At 20 weeks of age, *Esr1* expression was decreased (P=0.03), but *Pgr* was unaffected relative to controls (**Fig. 2O**, P=0.21).

*Ihh* is a progesterone-responsive gene and mediator of progesterone action for epithelial to stromal interactions in the mouse uterus (Lee et al., 2006; Takamoto et al., 2002). Expression of *Ihh* in mutants and controls was similar at all three timepoints (**Fig. 2Q**, P=0.31, P=0.29, P=0.07, respectively). *Foxa2* is a molecular marker of uterine glands (Besnard et al., 2004). *Foxa2* expression was similar between mutants and controls at 3, 8, and 20 weeks of age (P=0.49, P=0.24, P=0.29, respectively), but was lower in edematous mutant uteri at 8 weeks of age (**Fig. 2R**, P=0.04).

### Uterine pathologies in *Gnrhr^E90K^* homozygotes

At 8 weeks of age, the gross morphology of uteri from *Gnrhr*^*E90K*^ homozygotes appeared similar to controls except for swelling due to edema (**Fig. 3A, B**). By 20 weeks of age, we noticed dark focal regions within the uterine wall of the mutants that were visible under the stereo dissecting microscope (**Fig. 3C, D**). Histological analysis showed enlarged cavities peripheral to the uterine lumen associated with the glandular epithelium (**Fig. 2K, L**). In addition, there was squamous neoplasia, tubal neoplasia, and cribriform gland architecture, indicating glandular hyperplasia (**Fig. 3E, F**). Thus, by 20 weeks of age persistent estrous levels of estrogen and low progesterone led to uterine epithelial pathologies.

**Fig. 3.**
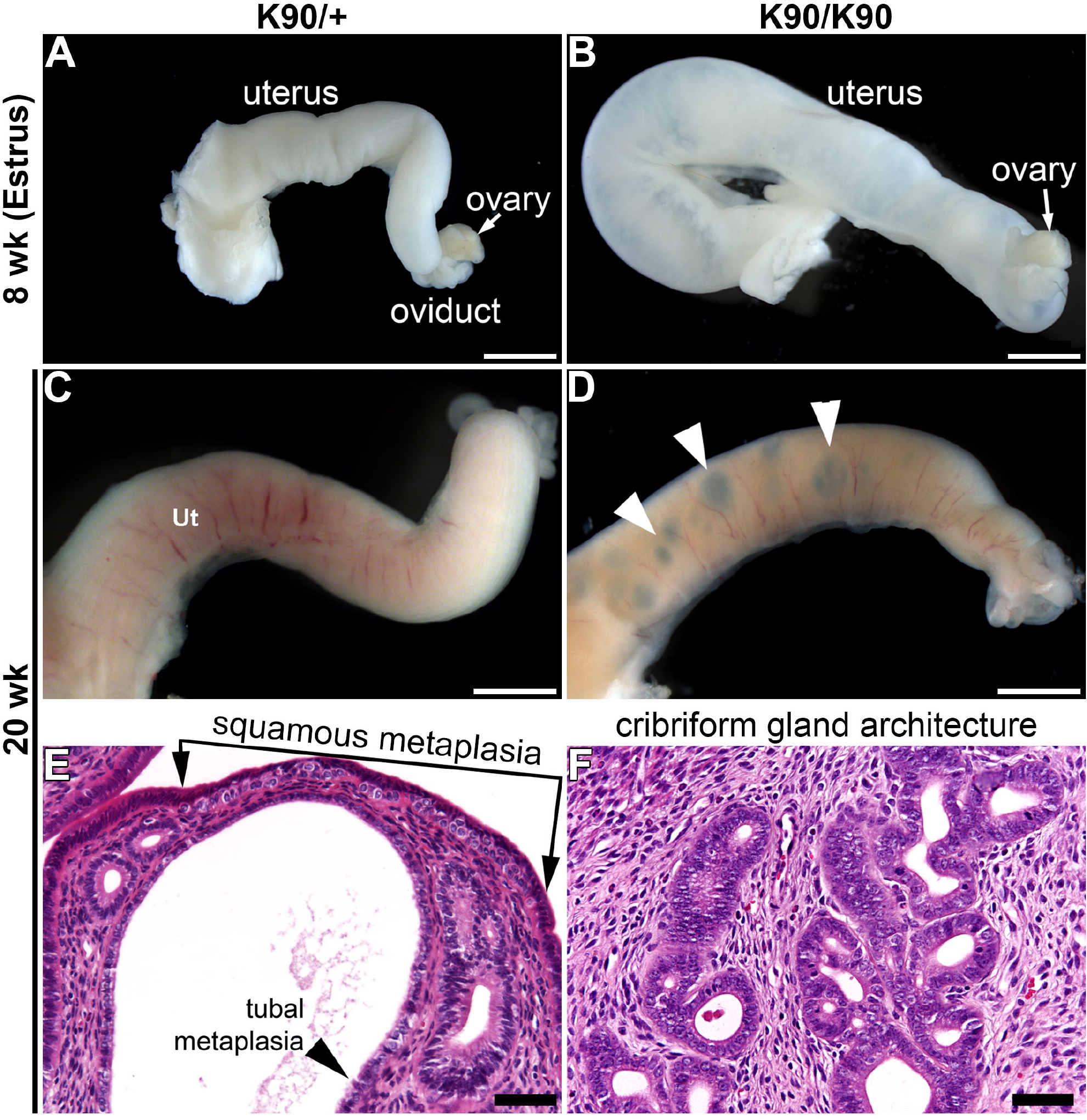
Uterine pathologies in *Gnrhr*^*E90K*^ homozygotes. (A-D) Dissecting stereomicroscopic images of uterine horns. (A, C) *Gnrhr*^*E90K*^ controls; (B, D) *Gnrhr*^*E90K*^ homozygous mutants. Focal abnormalities appeared by 20 week of age (white arrowheads in D). (E, F) H&E-stained sections of uteri at 20 weeks of age showing pathologies. A-D, scale bars = 2 mm; E,F 50 μm.

### Induced ovulation eliminates uterine edema, but does not rescue post-pubertal uterine gland defects in *Gnrhr^E90K^* homozygous females

To test whether repeated inductions of ovulation could rescue post-pubertal uterine gland development, *Gnrhr*^*E90K*^ homozygous females were treated with human chorionic gonadotropin (hCG) or vehicle every 4 days from 5 to 8 weeks of age. Administration of hCG induced ovulation and formation of CL in the mutants (**Fig. 4A, B**, **Supplemental Fig. 1A-C**). Uterine edema was observed in ~50% of the mutants treated with vehicle. In contrast, uterine edema was not observed in the hCG-treated mutants as shown by gross morphology (not shown), histology (**Fig. 4C, D**), and reduced uterine horn length (**Fig. 4E**, P=0.01). However, the repeated induced-ovulations did not rescue the post-pubertal uterine gland mutant phenotype (**Fig. 4F**, P=0.12). The lack of uterine edema in the hCG-treated mutants suggests that this treatment regimen was sufficient to alter hormone levels, but not to rescue adolescent uterine gland development.

**Fig. 4.**
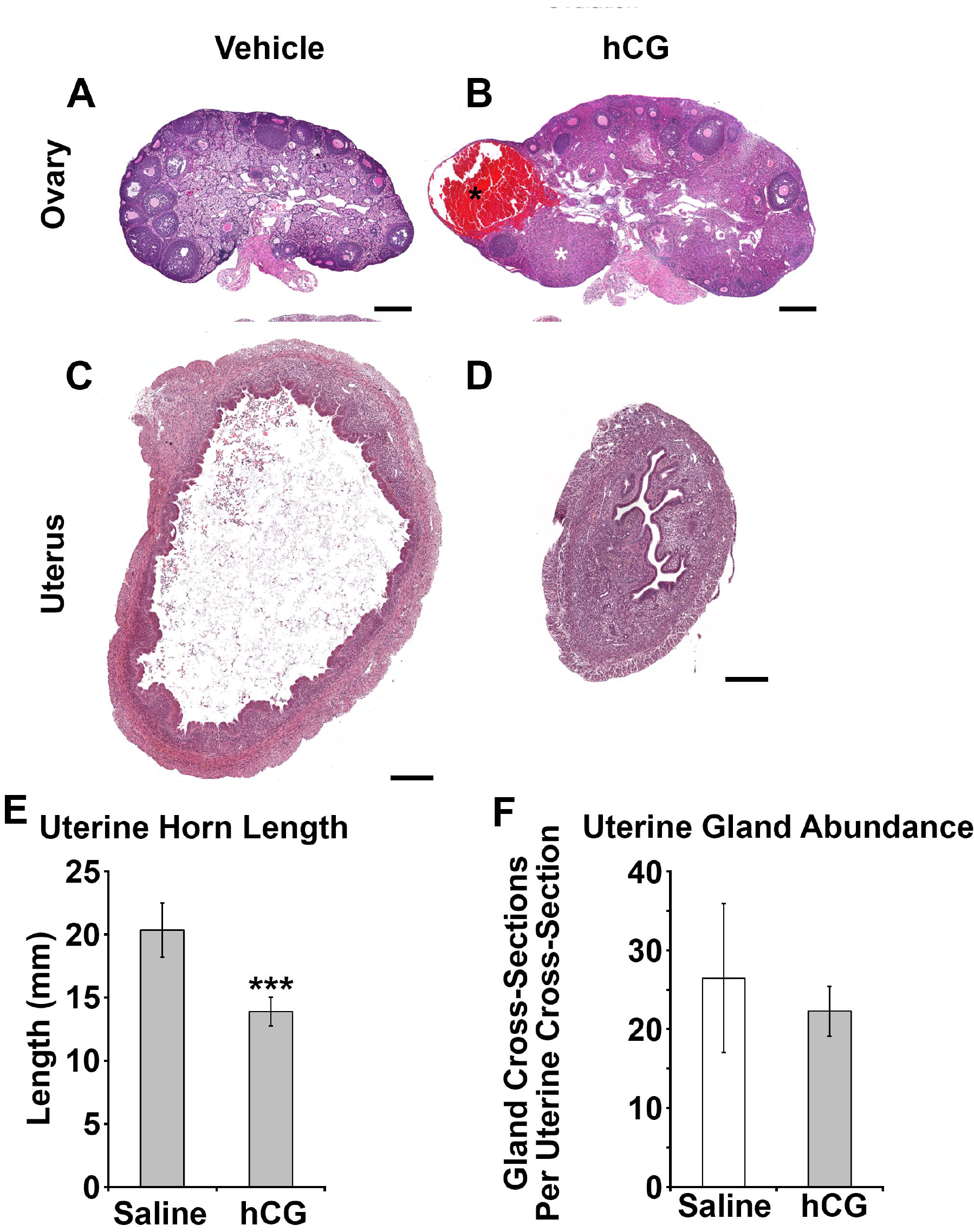
Induced ovulation eliminates uterine edema, but does not rescue uterine gland development in *Gnrhr*^*E90K*^ mice. (A,B) Administration of hCG induced ovulation as evident by development of corpora hemorrhagica (black asterisk) and corpora lutea (white asterisk) in *Gnrhr*^*E90K*^ homozygotes. (C,D) Administration of hCG restored uterine histoarchitecture in *Gnrhr*^*E90K*^ homozygotes. Scale bars = 250 μm. (E) Uterine horn length was reduced in mice that received hCG. (F) Uterine glandular tissue abundance was unaffected by hCG treatment. *** P < 0.01, Student’s *t*-test.

### Progesterone receptor knockout mice also have reduced post-pubertal uterine gland development

*Gnrhr*^*E90K*^ homozygous females fail to ovulate or form CL. In addition to estrous levels of estradiol, they have low levels of circulating progesterone. Thus, we hypothesized that the delayed post-pubertal gland development was due to insufficient progesterone signaling within the endometrium. To test this idea, we determined if post-pubertal gland development was also impaired in *Pgr* knockout (PRKO) mice. As previously reported (Lydon et al. 1995), the ovaries of PRKO females showed no evidence for ovulation (no CL were present in histological sections) (**Fig. 5A, B**). At 3 weeks of age, uterine histology was similar between PRKO females and controls (**Fig. 5C, D**). There was no difference in uterine horn length (**Fig. 5I**, P=0.13) or glandular tissue abundance between mutants and controls (**Fig. 5J**, P=0.40). By 8 weeks of age, uterine gland development was noticeably impaired (**Fig. 5E, F**), including reduced uterine horn length (**Fig. 5I**, P<0.005) and less glandular tissue (**Fig. 5J**, P<0.001). Similar to *Gnrhr*^*E90K*^ homozygotes, uterine gland development was comparable to controls by 20 weeks of age. However, PRKO uterine glands were dilated (**Fig. 5G, H**). These data suggest that progesterone signaling is important for uterine gland development during the post-pubertal period.

**Fig. 5.**
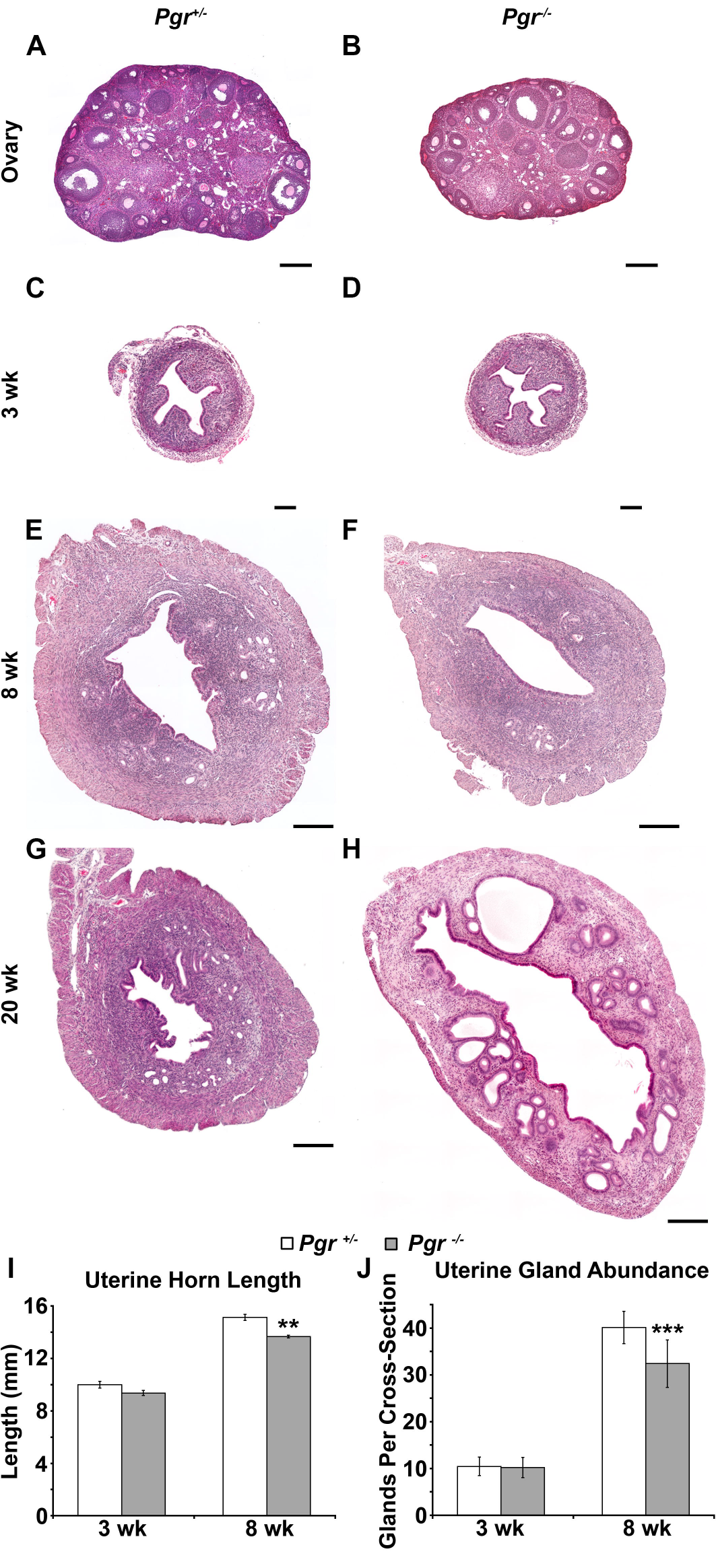
PRKO mice also display reduced uterine gland development at 8 weeks of age. (A, B) H&E-stained ovarian sections. Similar to *Gnrhr*^*E90K*^ homozygotes, PRKO mice exhibit ovulation failure. (C-H) H&E-stained uterine sections. The uteri of PRKO mice appeared morphologically similar at 3 weeks of age (C, D), but uterine glandular tissue was less abundant at 8 weeks of age (E, F). Similar to *Gnrhr*^*E90K*^ homozygotes, uterine glands were large and dilated at 20 weeks of age (G, H). A,B, scale bars = 250 Mm; C-H, scale bard = 200 μm. (I) Uterine horn length was reduced for PRKO females at 8 weeks of age. (J) Uterine glandular tissue abundance was reduced for PRKO females at 8 weeks of age. ** P<0.005, *** P < 0.001, Student’s *t*-test.

### Progesterone treatment does not rescue post-pubertal uterine gland development in *Gnrhr^E90K^* homozygous mice

Next, we tested the idea that low progesterone levels were responsible for delayed post-pubertal uterine gland development in *Gnrhr*^*E90K*^ homozygous mice. *Gnrhr*^*E90K*^ homozygous females were given vehicle or progesterone implants at 5 weeks of age and tissues were collected at 8 weeks of age (**Fig. 6A**). We confirmed elevated progesterone blood levels in the progesterone-treated group by radioimmunoassay (**Fig. 6B**, P<0.001). There was no effect of progesterone treatment on circulating levels of estradiol (P=0.16). Progesterone treatment did not affect ovarian histology as abundant antral follicles and no CL were observed in both vehicle and progesterone-treated mutants (**Fig. 6C, D**). Surprisingly, progesterone treatment did not rescue post-pubertal uterine gland development (**Fig. 6E, F**). Uterine horn length was reduced in the progesterone-treated group (**Fig. 6G**, P=0.01), perhaps because uterine edema was not observed. Uterine glandular tissue abundance was reduced by progesterone treatment (**Fig. 6H**, P<0.001). Thus, continual progesterone release relieved uterine edema, but negatively affected post-pubertal uterine gland development.

**Fig. 6.**
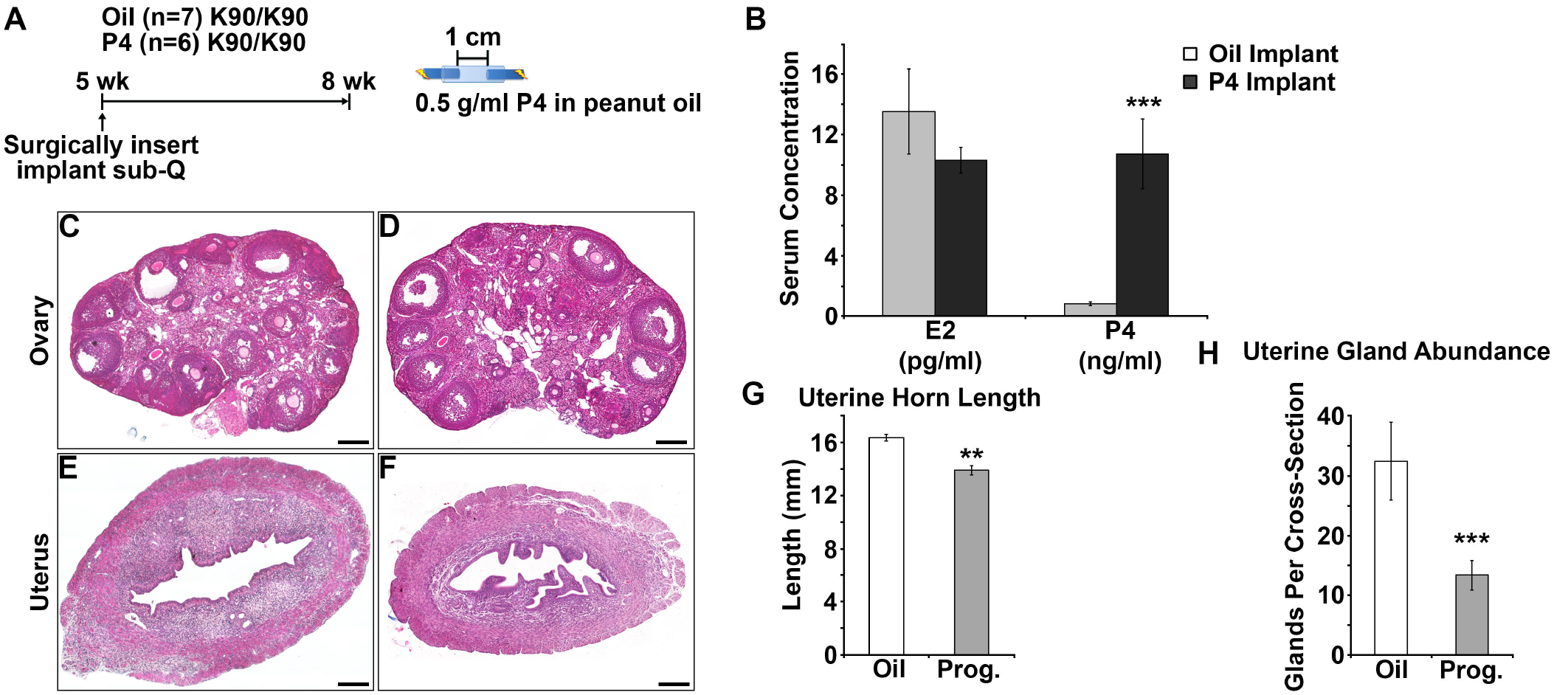
Progesterone supplement is unable to rescue uterine gland development in *Gnrhr*^*E90K*^ mice. (A) Experimental design. Slow release hormone implants were administered at 5 weeks of age and tissues harvested at 8 weeks of age. Only *Gnrhr*^*E90K*^ homozygotes were used for this study. (B) Administration of the progesterone implants increased serum progesterone levels, but did not affect estradiol levels. (C, D) Administration of progesterone did not affect the persistent antral follicle phenotype. (E, F) Overall uterine morphology was similar between groups; however, mice in the progesterone treated group appear to have less glandular tissue. (G) Uterine horn length was reduced in progesterone-treated mice. (H) Uterine glandular tissue abundance was reduced in progesterone-treated mice. Scale bars = 200 μm. ** P<0.05, *** P < 0.005, Student’s *t*-test.

## Discussion

The *Gnrhr*^*E90K*^ mouse model offers a unique opportunity to observe the consequences of reduced plasma membrane expression of GnRHR on the hypothalamic-pituitary-gonadal axis. Gonadotropes synthesize and release LH and FSH in response to the GnRHR ligand GnRH. GnRHR plasma membrane levels vary throughout the estrous cycle, peaking at proestrus in close correlation with the pre-ovulatory LH surge (Marian et al., 1981). At proestrus, GnRH release becomes less pulsatile and more sustained. *Gnrhr*^*E90K*^ homozygous females fail to produce an LH surge and therefore do not ovulate, resulting in an ovarian cycle that is arrested at the antral follicle stage. These observations support the idea that high levels of GnRHR plasma membrane expression are required to decode the switch from pulsatile to sustained GnRH. Thus, there is a close relationship between plasma membrane levels of GnRHR and gonadotropin release. Importantly, a threshold level of plasma membrane-associated GnRHR must be achieved to induce the LH surge.

The ovarian cycle of *Gnrhr*^*E90K*^ females is effectively arrested at the antral follicle stage due to ovulation failure. As a consequence, ovarian steroid hormones are not produced cyclically. We found continual production of estradiol at levels similar to those of an animal in estrus and very low progesterone. Low progesterone levels are a consequence of the inability to form CL. Thus, *Gnrhr*^*E90K*^ homozygous females exhibit a unique endocrine phenotype in which the estrous cycle is arrested. This phenotype has relevance to human polycystic ovarian syndrome (PCOS), which is also characterized by ovulation failure in the presence of multiple antral follicles (polycystic ovaries) (Bellver et al., 2017). The *Gnrhr*^*E90K*^ females exhibited neoplastic changes in their uterine epithelium by 20 weeks of age. Women with PCOS have an increased risk of developing endometrial cancer (Dumesic and Lobo, 2013). Importantly, our data indicate that PCOS is likely to be detrimental to uterine gland growth and maintenance of pregnancy. Thus, even if ovulation is restored, the ability of the uterus to support peri-implantation growth and development might be compromised. This could account for the increased prevalence of low birth weight babies born to women with PCOS (Sir-Petermann et al., 2005).

Uterine gland development occurs in two waves. The neonatal wave is considered to be ovarian hormone independent, whereas the post-pubertal (adolescent) wave is hormone dependent. In *Gnrhr*^*E90K*^ homozygous females, the hormone-independent neonatal wave proceeded similar to controls, but the second wave of development was impaired. These results support the idea that the neonatal wave is hormone-independent and expand our understanding of the post-pubertal wave by illustrating the importance of cyclic, rather than continual, hormone production for proper uterine gland growth and development.

Not only was uterine gland development impaired, but disease-related phenotypes manifested, including edema and endometrial hyperplasia. Interestingly, edema appeared in about 50% of *Gnrhr*^*E90K*^ homozygotes at 8 weeks of age, but in none of the E90K mice at 20 weeks of age. Thus, absence of edema correlated with appearance of neoplasms. Perhaps persistent high levels of estradiol first stimulated uterine edema and then, with time, promoted neoplasia. In one mouse model, estrogen treatment resulted in uterine edema (Wu et al., 2015). These observations illustrate the negative consequence of persistent estrous levels of estradiol in the absence of progesterone.

PRKO mice shared reproductive organ phenotypes with *Gnrhr*^*E90K*^, including reduced uterine gland development and uterine edema; however to a lesser extent. The PRKO uterine phenotype may be the result of similar endocrine alterations, including a lack of progesterone signaling and elevated estradiol. These data support the idea that progesterone signaling in the uterus counteracts the effects of estradiol on uterine edema.

We tested whether progesterone treatment could rescue the uterine defects in the *Gnrhr*^*E90K*^ mutants by administering slow-release progesterone implants between 5 and 8 weeks of age. Progesterone treatment eliminated uterine edema, but did not rescue the second wave of uterine gland development. Thus, progesterone counteracted the effects of estradiol to suppress edema, but was not sufficient to rescue post-pubertal uterine gland development. This may be due to the continual release of progesterone by the implant. Progesterone levels rise and fall during the estrous cycle in an inverse relationship to estradiol. This suggests that cyclic—not continuous— hormone secretion is necessary for correct uterine gland growth during adolescence.

We attempted to rescue the uterine defects of the *Gnrhr*^*E90K*^ homozygotes by administering hCG every 4 days to restore cyclicity. As expected, this protocol induced ovulation and luteinization in the mutants. Uterine edema was eliminated, but the second wave gland development was not rescued. These observations indicate that the uterine edema is fueled by chronic estradiol secretion from the persistent antral follicles. Furthermore, these data suggest that uterine gland growth is very sensitive to the endogenous (optimal) pattern of hormone secretion, which cannot be fully replicated by hCG injection.

Both progesterone implants and hCG injections prevented uterine edema in *Gnrhr*^*E90K*^ homozygous females. The progesterone treatment likely counteracted the effects of chronic estradiol and the hCG injections induced ovulation, eliminating the persistent antral follicles. Thus, both experiments show that persistent antral follicles are the cause of the uterine edema in these mice. To our knowledge uterine edema has either not been reported or not studied in women with PCOS; however, fluid retention and general edema are common. Women with PCOS are typically treated with progesterone or cyclic birth control to simulate an estrous cycle. Our data suggest that these hormone treatments may be sufficient to overcome possible uterine edema, but not to rescue uterine gland numbers in women with PCOS.

In conclusion, post-pubertal uterine gland growth and homeostasis requires proper regulation of cyclic ovarian hormone secretion. In our mouse model, polycystic ovaries had detrimental effects on post-pubertal uterine gland growth, which could impair peri-implantation conceptus development. Finally, the absence of cyclic steroid hormone production can lead to epithelial pathologies of the uterus, including neoplasia.

## Acknowledgment

We are grateful to Franco DeMayo (National Institute of Environmental Health Sciences) for generously providing *Pgr*^*tm2(cre)Lyd*^ mice. We dedicate these studies to Dr. P. Michael Conn.

## Supplemental Figure Legends

**Suppl. Fig. S1. A single injection of hCG induces ovulation in *Grnhr*^*E90K*^ homozygous mice.** (A, B) H&E-stained ovarian sections. Corpora lutea (CL) were absent from controls (A), but abundant in the ovaries of mice that received hCG (black asterisks) (B). (C) Average number of CL per female (both ovaries) (± SEM). Scale bars = 200 μm. hCG, human chorionic gonadotropin.

